# Dose-Dependent Induction Of CPP Or CPA By Intra-pVTA Ethanol: Role Of Mu Opioid Receptors And Effects On NMDA Receptors

**DOI:** 10.1101/2020.01.07.897702

**Authors:** Yolanda Campos-Jurado, Lucía Martí-Prats, Jose A. Morón, Ana Polache, Luis Granero, Lucía Hipólito

## Abstract

The neurobiological mechanisms underlying alcohol motivational properties are still not fully understood, however, the mu-opioid receptors (MORs) have been evidenced as central elements in the manifestation of the alcohol reinforcing properties. Drug-associated environmental stimuli can trigger alcohol relapse and promote alcohol consumption whereby N-methyl-D-aspartate (NMDA) receptors play a pivotal role. Here we sought to demonstrate, for the first time, that ethanol induces conditioned place preference or aversion (CPP or CPA) when administered locally into the ventral tegmental area (VTA) and the associated role of MORs. We further analyzed the changes in the expression and mRNA levels of GluN1 and GluN2A subunits in designated brain areas. The expression of CPP or CPA was characterized following intra-VTA ethanol administration and we showed that either reinforcing (CPP) or aversive (CPA) properties are dependent on the dose administered (ranging here from 35 to 300 nmol). Furthermore, the critical contribution of local MORs in the acquisition of CPP was revealed by a selective antagonist, namely β-Funaltrexamine. Finally, modifications of the expression of NMDA receptor subunits in the Nucleus Accumbens (NAc) and Hippocampus after ethanol-induced CPP were analyzed at the proteomic and transcriptomic levels by western blot and *In Situ* Hybridation RNAscope techniques, respectively. Results showed that the mRNA levels of GluN2A but not GluN1 in NAc are higher after ethanol CPP. These novel results pave the way for further characterisation of the mechanisms by which ethanol motivational properties are associated with learned environmental cues.

## 1. Introduction

The neurobiological mechanisms underlying alcohol motivational properties are still not fully understood. Although it is well established that drug-associated environmental stimuli can trigger alcohol relapse and promote alcohol consumption (Bossert et al., 2013; Crombag et al., 2008), the studies aimed to demonstrate other context-ethanol associations have yielded contradictory results.

One of the paradigms widely accepted to approach these context-drug associations in preclinical research is the conditioned place preference (CPP). Rats can develop a preference for an environment due to the association of this context with the reinforcing properties of the drug through the activation of the mesocorticolimbic system (MCLS). This has been demonstrated for the majority of drugs of abuse. However, the literature has yielded paradoxical results when alcohol is administered, due to its complex pharmacology. Some studies showed that systemic administration of low to medium ethanol doses (from 0.25 to 1 mg/Kg) induced CPP in rats, whereas in other studies, rats did not show any preference for the ethanol-paired compartment (Bahi and Dreyer, 2013; Peana et al., 2008; Zarrindast et al., 2010). Interestingly, when high doses of ethanol (2 g/kg) are used, rats appear to develop an aversion to the drug-associated context (Becker et al., 2006). The differences in experimental design could potentially contribute to the disparity of behavioral responses, thus, the number of conditioning sessions, the route of administration or the dose of ethanol administered have shown to be essential in the expression of CPP. Furthermore, our previous work revealed that the balance between the ethanol molecule and its metabolic products in the MCLS could determinate the reinforcing properties of this drug (Martí-Prats et al., 2015, 2013).

According to this hypothesis, enzymatic systems responsible for the biotransformation of ethanol in the brain play a crucial role on the motivational properties of ethanol. Depending on the dose administered and the associated saturation of the enzymes, we should see either an activation, an inhibition, or no changes in the activity of the dopaminergic (DA) neurons of the MCLS. The ethanol molecule itself (the non-metabolized fraction) may trigger inhibition of the DA neurons within the ventral tegmental area (VTA) by increasing gamma-amino butyric acid (GABA) release from presynaptic terminals, presumably through the GABA A receptors (Xiao et al., 2009). Our previous studies have shown that this non-metabolized fraction is responsible for the depressant effects of ethanol after its administration into the VTA, elucidated as a decreased locomotor activity (Martí-Prats et al., 2013). This phenomenon has been shown not only at the behavioral level but also at the electrophysiological level (Theile et al., 2011, 2009, 2008). Furthermore, increasing evidence support the idea that salsolinol, a metabolic derivative of ethanol, would be responsible for the excitation of the VTA DA neurons (Hipólito et al., 2011; Martí-Prats et al., 2015; Melis et al., 2015; Xie et al., 2012). Salsolinol activates the mu opioid receptors (MORs) expressed in VTA GABA neurons that tonically inhibit the activity of VTA DA neurons and, as a consequence, reduces GABA release, disinhibiting DA neurons (Xie et al., 2012). This mechanism is similar to the one proposed by Johnson and North explaining VTA DA activation by opioids (Johnson and North, 1992). Evidences supporting this hypothesis have demonstrated that salsolinol, acting as an agonist of the MORs (Berríos-Cárcamo et al., 2016; Fertel et al., 1980; Lucchi et al., 1982), consequently triggers increased DA levels in the nucleus accumbens (NAc) and produces opioid-like effects (Hipólito et al., 2010; Melis et al., 2015; Xie et al., 2012). Additionally, rats readily self-administer salsolinol, as well as ethanol, when infused in the posterior but not anterior VTA, demonstrating the reinforcing properties of this ethanol metabolite (Rodd et al., 2008, 2005) (for further review in salsolinol effects see (Hipólito et al., 2012). Within this framework, we hypothesized that administration of low to medium ethanol doses (associated in previous experiments with an increased activity of VTA DA neurons), when paired to a specific environment, would induce the development of CPP. However, administration of higher ethanol doses (causing GABA release and DA neurons inhibition (Jhou et al., 2009; Tan et al., 2012; Ungless et al., 2004)) would trigger conditioned place aversion (CPA).

Repeated activation of VTA DA neurons by exposure to psychostimulants and opiates triggers glutamatergic adaptation both at pre-and postsynaptic levels in the NAc (reviewed in (Hearing et al., 2018)) and beyond, e.g. the Hippocampus (Fakira et al., 2014; Portugal et al., 2014). Similarly, adaptations in the glutamatergic synapses have also been described following repeated administration of ethanol (Flatscher-Bader et al., 2008; Kemppainen et al., 2010; Wills et al., 2012; Xiao et al., 2009). In fact, the N-methyl-D-aspartate glutamate receptor (NMDAR), which is a regulator of drug-induced plasticity in excitatory synapses, plays a key role in the environment-drug associations, especially when the drug pharmacology is dependent on MORs (Fakira et al., 2014; Portugal et al., 2014; Sikora et al., 2016). In fact, antagonizing NMDAR blocks ethanol-induced CPP expression (Gremel and Cunningham, 2009) and NMDA 2A receptor (GluN2A) KO mice exhibit a loss of ethanol-induced CPP (Boyce-Rustay and Holmes, 2006). However, few studies have investigated how association between an environment and the motivational properties of ethanol could modify the expression of the NMDAR in key brain areas such as NAc and Hippocampus.

Considering all this evidence, in the present study we aimed to characterize, for the first time, the ability of ethanol to induce CPP or CPA when administered directly into the posterior VTA (pVTA), emphasizing on the role of the MORs in this phenomenon. In addition, we analyzed the modifications in the expression of NMDA 1 (GluN1) and GluN2A subunits of the NMDAR in the NAc and Hippocampus following the induction of CPP or CPA. We found that ethanol dose-dependently induces CPP or CPA when administered locally in the pVTA, triggering changes in the GluN2A mRNA levels in the NAc without affecting its expression in the Hippocampus. Moreover, and in accordance with the previous published work, we showed here that MORs play a critical role in ethanol-induced CPP.

## 2. Methods

### 2.1 Animals

Male albino Wistar rats (300-340 g at the time of surgery) were housed in plastic cages (48 x 38 x 21 cm^3^) in groups of four to six in a humidity and temperature controlled environment (22°C), under a 12:12-h light/dark cycle (on 08:00, off 20:00) with ad libitum access to food and water. All the procedures were carried out in strict accordance with the EEC Council Directive 86/609, Spanish laws (RD 53/2013) and animal protection policies. Experiments were approved by the Animal Care Committee of the University of Valencia and authorized by the Regional Government.

### 2.2 Drugs and chemicals

Ethanol was purchased from Scharlau (Madrid, Spain). β-funaltrexamine (β-FNA; an irreversible antagonist of the MORs) was obtained from Tocris (Bristol, UK). Stock solutions of β-FNA were prepared by dissolving the compound in the correct volume of distilled water to obtain 13.6 mM concentration of β-FNA. Aliquots of these solutions were then kept frozen at −20°C until use. Prior to use, aliquots of the stock solutions were conveniently diluted with artificial cerebrospinal fluid (aCSF) solution to the appropriate concentration (8.3 mM) (Sánchez-Catalán et al., 2009). Ethanol was also freshly dissolved in aCSF solution (from 87.5 to 750 mM) prior to intra-pVTA administration. The aCSF solution was prepared and stored at −20°C. Its composition was: 120.0 mM NaCl, 4.8 mM KCl, 1.2 mM KH_2_PO_4_, 1.2 mM MgSO_4_, 25.0 mM NaHCO_3_, 1.2 mM CaCl_2_, 100 mM d-glucose, and 0.2 mM ascorbate, pH adjusted at 6.5 (Hipólito et al., 2010).

All the other reagents used were of the highest commercially available grade.

### 2.3 Surgery and post-surgical care

Rats were anesthetized with ketamine/xylazine (80 mg/kg of ketamine and 10 mg/ kg of xylazine), intraperitoneal (i.p.) and placed in a stereotaxic apparatus (Stoelting, USA). An incision (8-10 mm) was made in the skin over the skull and the wound margin was infiltrated with lidocaine (3%). Four holes were drilled: two for the skull screws and the other two for the guide cannulae (Plastics One, USA). Each animal was implanted bilaterally with two 28-gauge guide cannulae aimed at 1.0 mm above the pVTA. The coordinates, taken from the bregma and skull surface (Paxinos and Watson, 2007) were as follows: A/P −6.0 mm; L ±1.9 mm; D/V −7.8 mm (Martí-Prats et al., 2013). Cannulae were angled toward the midline at 10° from the vertical (all the measurements in the dorsal-ventral plane refer to distances along the track at 10° from the vertical). Cannulae assemblies were secured with dental cement. A stainless-steel stylet (33-gauge), extending 1.0 mm beyond the tip of the guide cannula, was inserted at the time of surgery and removed at the time of testing. Following surgery, rats were housed in individual rectangular plastic cages (47 x 22 x 15 cm^3^, placed side by side in order to prevent any potential influence of chronic stress on performance due to isolation) with free access to food and water.

### 2.4 Drug microinjection procedures

All the intra-pVTA drug microinjections were carried out with 33-gauge stainless steel injectors, protruding 1.0 mm below the tip of the guide cannulae. Injectors were attached to a 25 µL Hamilton syringe by PE-10 tubing. Microinjections were performed using a syringe pump (Kd Scientific) programmed to deliver a total volume of 200 nL in 20 s (flow rate of 0.6 mL/min), excepted for the β-FNA injections whereby the syringe pump was programmed to deliver a total volume of 300 nL in 2 min (flow rate of 0.15 µL/min). Following the infusion, the injectors remained in place for 1.5 min to allow the diffusion of the drugs, and was subsequently removed. All the injections were carried out in the experimental room and animals were placed in the CPP box immediately after injection (maximum latency time of 10 seconds).

### 2.5 Behavioral experiments

#### 2.5.1 CPP general procedure

CPP test was conducted in an unbiased manner whereby the two-compartments of the experimental box where separated by a barrier with a central door. The two compartments differed by the wall color: black and white vertical stripes (vertical compartment) and black and white horizontal stripes (horizontal compartment). Prior to and after the surgery, animals were handled every day for 4 days. Following surgery recovery, animals were exposed to the CPP box for 5 min in order to habituate them to the apparatus. The day prior to the conditioning, animal natural preference for one compartment was tested during 15 min (Pretest). During Conditioning, rats received bilateral intra-pVTA infusions prior to be immediately placed in the appropriate compartment. The conditioning phase consisted in 8 sessions (2 sessions/day: morning session and afternoon session, with a counterbalanced assignation of treatment and an interval of at least 5 hours between sessions) of 30 min distributed over 4 days. Animals were randomly assigned to the experimental or control group and the exposure to conditioning compartments was counterbalanced in both groups. After the last conditioning session, each animal was tested for its place preference (Test); the rat was placed in the open door of the barrier, and the time spent in each compartment was recorded over 15 min. Place preference scores were calculated as test minus pre-test time spent (in seconds) on the ethanol-paired compartment by visualizing the video recording of the pre-test and test sessions. Analysis of the behaviour was made to ensure a blind design by coding the name of the videos. Between 1.5 and 2 hours after the beginning of the test, rats were anaesthetized using isoflurane and brains were harvested and quickly frozen for further western blot or *In Situ* hybridization analyses and for the assessment of the cannulae placements (Figure. 1A).

**Figure 1.**
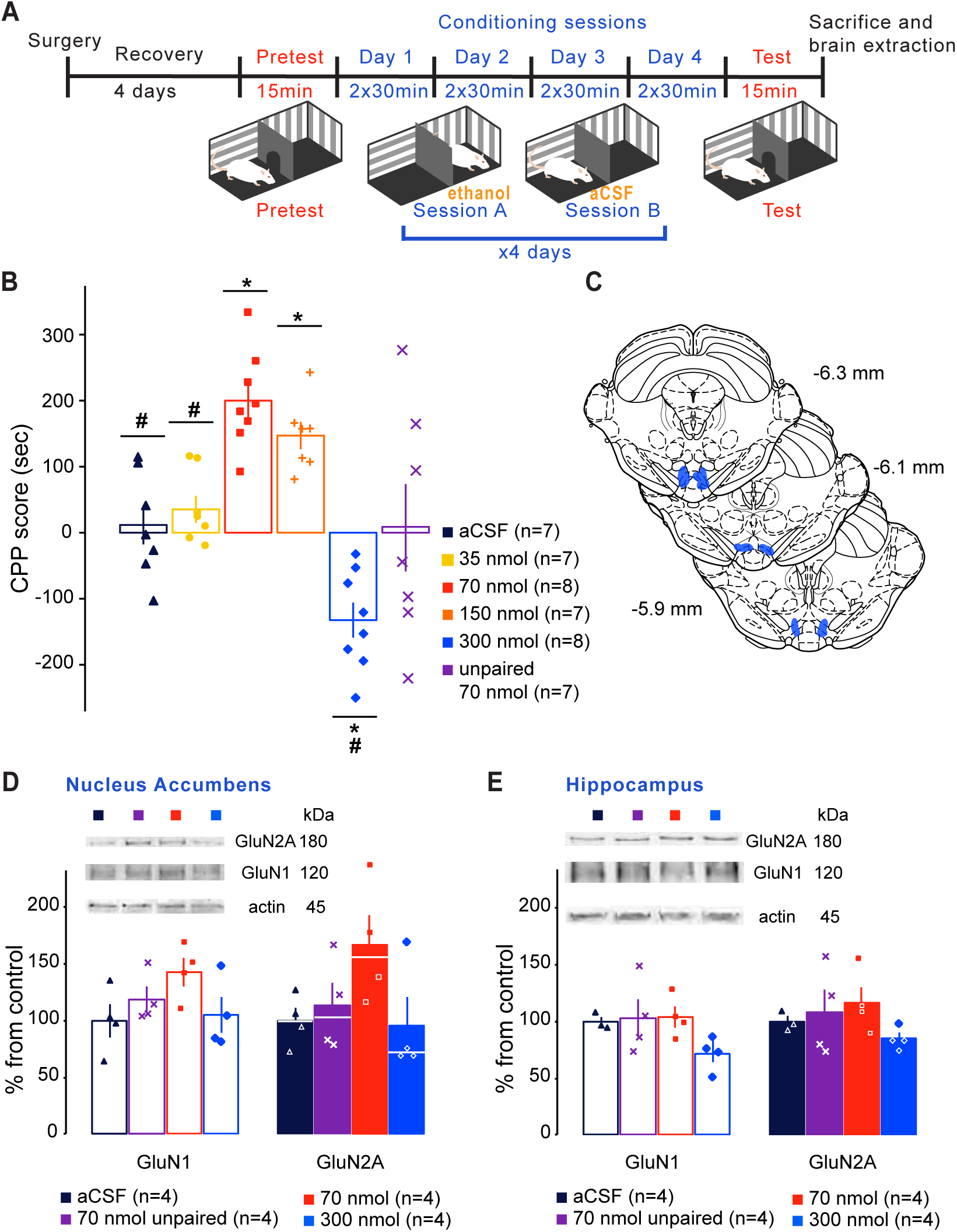
Ethanol intra-pVTA induces either CPP or CPA in a dose-dependent manner. ***A***, Schematic of the experimental design. ***B***, Place preference elicited by the administration of intra-pVTA ethanol (35, 70, 150 or 300 nmol). Data are represented as mean ± SEM of the preference score (test minus pretest time spent in ethanol-paired compartment). * denotes significant differences relative to the control group (aCSF). # denotes significant differences relative to the 70 nmol treated group. (* p<0.05, ** p<0.01, # p<0.01, ## p<0.001; Brown-Forsythe followed by Games-Howell test). ***C***, Diagram of brain coronal sections indicating in blue the area of all microinjections in pVTA. ***D***, NMDA subunit expression in NAc after ethanol induced CPP or CPA. Data are represented as mean ± SEM of the percentage from control group (aCSF). The withe line represents the median for the non-normal distributed data. ***E***, NMDA subunit expression in hippocampus after ethanol induced CPP or CPA. Data are represented as mean ± SEM of the percentage from control group (aCSF).

#### 2.5.2 Experiment 1: Intra-pVTA ethanol dose response CPP

Following the previously described procedure, 37 rats (n=7-8 per group) were randomly assigned to one of the four experimental groups receiving different doses of ethanol infused bilaterally intra-pVTA: 35 nmol, 70 nmol, 150 nmol and 300 nmol. Doses were selected from our previous published experiments covering from low to high doses (Martí-Prats et al., 2013; Sánchez-Catalán et al., 2009). Control group animals (n=7) received 8 infusions of the equivalent volume of aCSF while the other groups received ethanol or aCSF on alternate sessions. As a control for the effect of ethanol administration by itself, an additional group, the so-called ethanol-unpaired control group, was designed: in animals belonging to this group, the association of ethanol administration (70 nmol) with the compartment was alternated between days (i.e., horizontal compartment on day 1, vertical compartment on day 2 and so on) (Figure 1A).

#### 2.5.3 Experiment 2: MORs role in ethanol CPP acquisition

In this study, 29 animals received two bilateral intra-pVTA pre-treatments with 2.5 nmol of the MORs selective antagonist β-FNA or aCSF 24h before the first and third conditioning days. The β-FNA is an irreversible antagonist and, therefore, is able to block the MORs during 2 consecutive days following its administration as previously reported (Ward et al., 2003; Sánchez-Catalán et al., 2009). Then, during conditioning, rats received either 70 nmol of ethanol and aCSF (experimental group) or only aCSF (control group) (Figure 6A). Therefore, the different groups in this experiment were: aCSF + aCSF, aCSF + ethanol 70nmol, β-FNA + aCSF and β-FNA + ethanol 70nmol (Figure 6A).

### 2.6 Western-blot experiments

Serial coronal sections (40 µm thickness) containing the areas of interest were obtained using a cryostat. The A/P stereotaxic coordinates of each area (NAc and Hippocampus) were determined according to the atlas of Paxinos and Watson (Paxinos and Watson, 2007). Samples were obtained by punching a portion from an approximately 1-mm-thick coronal slice that included the areas of interest. Tissues were then homogenized in a 0.25 mM sucrose buffer containing 1 mM MgCl_2_ and protease inhibitors. The homogenates were then centrifuged at 13,200*g* for 10 min at 4°C to exclude large cells debris. The supernatants were used as total protein sample. Protein concentrations were measured using the Bio-Rad protein assay kit (Bio-Rad, Madrid, Spain).

Proteins were separated on SDS–polyacrylamide gel electrophoresis (PAGE) gels (4.5% acrylamide stacking gel and 10% acrylamide resolving gel) and were then transferred to 0.45 µm nitrocellulose membranes (Bio-Rad) using a semi-dry transfer system (Trans-Blot Turbo, Bio-Rad) for 30 minutes at 25V. Membranes were then blocked in 5% non-fat dried milk in TBS-Tween-20 (TBS-T) 0.1% (20mM Tris and 500mM NaCl pH 7.5) and incubated overnight at 4°C with the primary antibody: anti-GluN1 (1:1000; Merck KGaA, Darmstadt, Germany) and anti-GluN2A monoclonal antibody (1:1000; Merck KGaA, Darmstadt, Germany). After three washes in TBS-T, the blots were incubated for 1 hour with horseradish peroxidase– conjugated secondary antibody (1:1000; Bio-Rad, USA). Finally, blots were developed using the enhanced chemiluminescence system (ECL Plus, Amersham Bioscience, Little Chalfont, UK) according to the manufacturer’s protocol. Digital images of the immunoblots were obtained in a ChemiDoc imaging (BioRad, USA) system and further analysed using Image J software. The intensity of the bands at 170 and 105 kDa for GluN1 and GluN2A was expressed as arbitrary units and normalized by actin expression. Actin expression was analysed in the same blots by incubating with actin antibody (1:1000; Thermofisher, Ca, USA) followed by the incubation with the appropriate secondary antibody (1:1000 anti-IGgmouse; Invitrogen, Ca, USA) and developed with enhanced chemiluminescence system (Clarity Max ECL, BioRad, Cal, USA). The protein levels of the experimental groups (aCSF, ethanol 70 and 300 nmol and unpaired 70 nmol for the dose-response experiment; and aCSF + aCSF and β-FNA + ethanol 70nmol for the MOR blockade experiment) were determined by setting the control group (aCSF conditioned animals) to 100% and calculating the respective percentages, expressed as mean or median ± SEM.

### 2.7 mRNA *In Situ* hybridization Assay

Levels of GluR1 and GluR2A mRNA expression were examined by *in situ* hybridization using the RNAscope® technique (Advanced Cell Diagnostics, Hayward, CA, USA). Tissue was obtained from a new cohort which underwent the CPP general procedure divided in two groups: 70 nmol (n=6) and control (n=4). The *in situ* hybridisation was then carried out following the manufacturer guidelines, i.e. RNAscope® Sample Preparation and Pre-treatment Guide for Fresh Frozen Tissue, Part 1 (Document No. 320513-USM) and RNAscope® Fluorescent Multiplex Kit User Manual Part 2 (Document No. 320293-USM). Briefly, brains were cut on a cryostat into 15 µm thickness sections and collected on slides. The slides were then prepared by fixing and dehydrating the tissue and protease IV was applied. The hybridization and amplification were performed by using the Rn-Grin-C3 (317021-C3) and Rn-Grin2a (414621) probes (Advanced Cell Diagnostics, Hayward, CA, USA). Sections were then covered with coverslips and prepared for confocal microscopy assessment (Leica Biosystems, Germany). Images of the NAc and the Hippocampus for hemisphere were obtained with a 40xobjective. The quantification was carried out from combining images composed by 110 or 165 images of the NAc and Hippocampus, respectively (composition of 10 x 11 images of the NAc and 15 x 11 images of the Hippocampus), to cover all the areas of interest. Finally, quantification of the mRNA levels was conducted with the software FIJI following the RNAscope® Fluorescent Quantification Guidelines. The pictures files were coded by an investigator blind to the experimental design to guarantee unbiased analysis. Briefly, we counted the total number of cells and the average intensity per single dot was calculated for each merging file by measuring at least 20 single signal dots. The total intensity and average intensity per single dots were then used to calculate the total number of dots. Finally, data were expressed as the average number of dots per cell. Since the infusions were bilateral in all the experiments, the two-hemisphere data were averaged.

### 2.8 Histology

40 µm-thick coronal sections of the pVTA area were obtained using a cryostat. Sections were used for the assessment of the cannulae placements of all the experiments presented in this study. Sections were then stained according to a standard cresyl violet protocol. Then, the placement of the cannula tip was carefully examined by a researcher who was blind to the experimental condition of the animals, using optical microscopy (see fig 1C, 2C and 3C).

**Figure 2.**
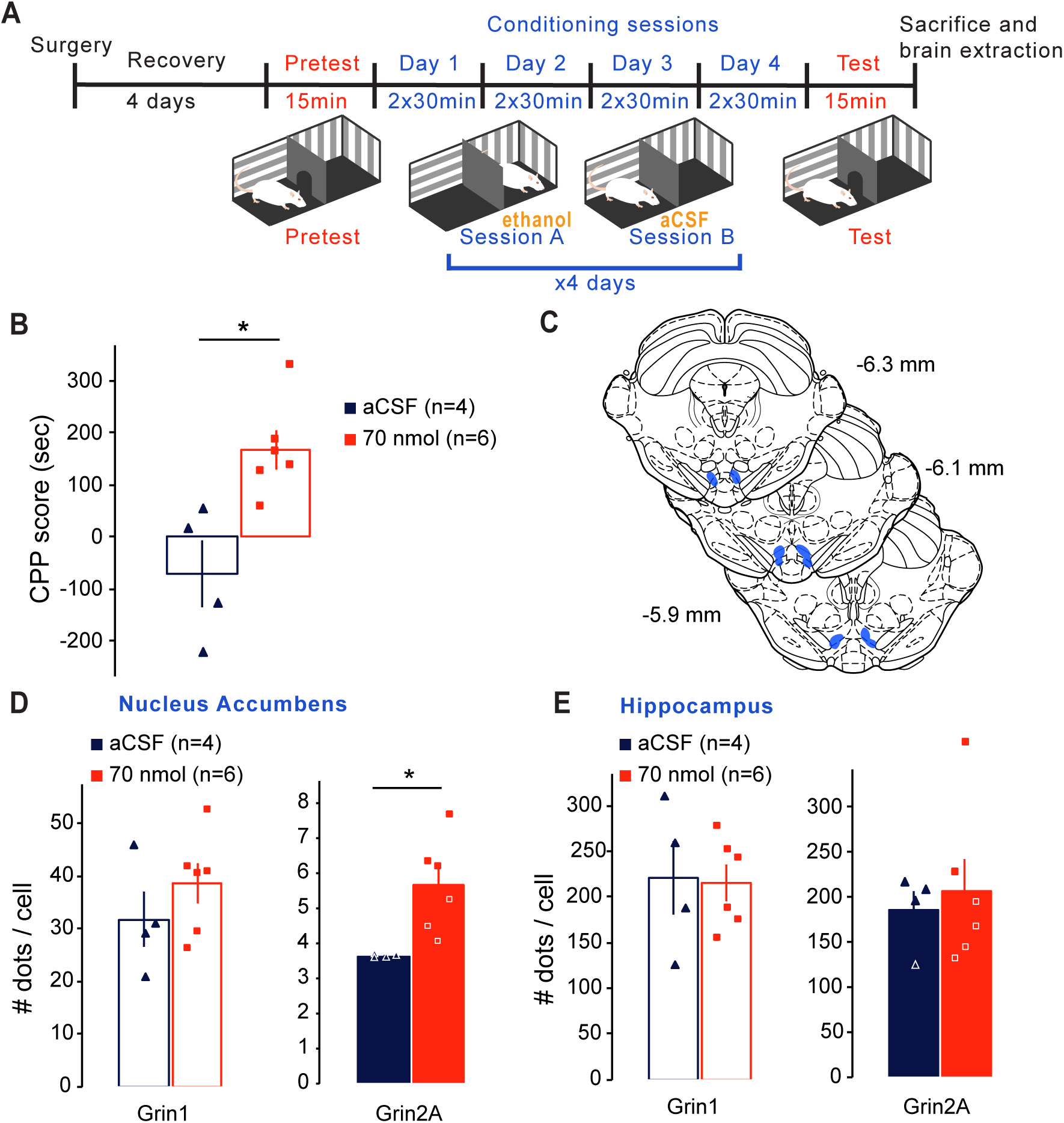
Ethanol place preferences is associated with the expression of mRNA from GluN2A but not from GluN1 subunit in the NAc. ***A***, Schematic of the experimental design. ***B***, Place preference elicited by the administration of 70 nmol of ethanol in the pVTA. Data are mean ± SEM represented as preference score (test minus pretest time spent in ethanol-paired compartment). * denotes significant differences between means (p<0.01, t-Test). ***C***, Diagram of brain coronal sections indicating in blue the area of all microinjections in pVTA. ***D***, Expression of mRNA from NMDA subunits GluN1 and GluN2A in NAc after ethanol induced CPP. Data are mean ± SEM represented as number of dots (mRNA molecules) per cell. * denotes significant differences between means (p<0.05). ***E***, Expression of mRNA from NMDA subunits GluN1 and GluN2A in hippocampus after ethanol induced CPP. Data are mean ± SEM represented as number of dots (mRNA molecules) per cell.

**Figure 3.**
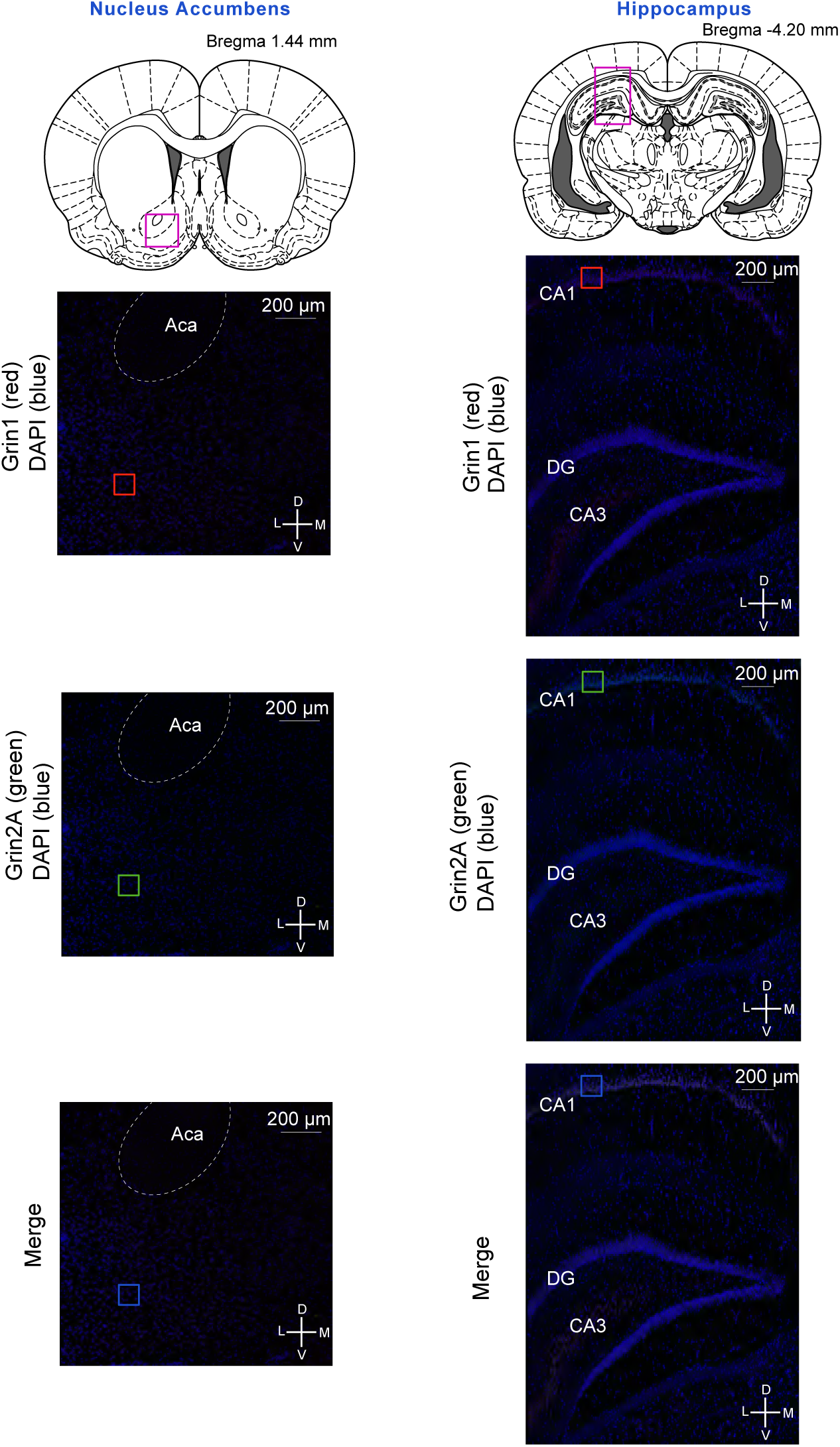
Representative pictures of the compilation files obtained in the mRNA *In Situ* Hybridization Assay used for the quantification of mRNA from GluN1 and GluN2A subunits in NAc (left) and hippocampus (right) after ethanol induced CPP. Squares delimit the area of the amplified pictures shown in Figure 4 for the NAc and in Figure 5 for the hippocampus. Abbreviatures: *Aca*, anterior commissure; *CA3*, field CA3 of hippocampus; *DG*, dentate gyrus; *CA1*, field CA1 of hippocampus.

### 2.9 Statistical methods

All data were expressed as Mean or Median ± SEM when appropriate.

Data from the experiments were analyzed by performing ANOVAs or T-tests, depending on the number of groups included in the analysis. In all cases, normality was tested before performing ANOVAs or T-tests. When the Shapiro-Wilk test revealed a non-normal distribution, differences between groups were analyzed using the Kruskal-Wallis non-parametric test and Bonferroni adjustment for pairwise comparisons. When normally distributed, data were analyzed using one-way ANOVA, followed by Tukey’s test. Homogeneity of variance was also tested before the ANOVA. When the assumption of the homogeneity of variances was violated, data were analyzed using Brown-Forsythe test of equality of means, followed by Games-Howell test.

Statistical analyses were performed with IBM SPSS statistics 24 software.

## 3. Results and statistical analyses

### Ethanol intra-pVTA induces either CPP or CPA in a dose-dependent manner

The mean values of the preference score, calculated by subtracting the time spent in the drug-paired side during the Pre-test session from the time spent in the drug-paired side during the Test session, for each intra-pVTA ethanol dose group are shown in Figure 1B. The homogeneity of variance was violated, thus the Brown-Forsythe adjustment was performed. Results showed that ethanol significantly modifies the time spent in the drug-paired compartment during the Test session compared to the Pre-test (p< 0.001). Both the groups administered with 70 and 150 nmol of ethanol showed a significant increase in the preference score for the drug-paired compartment as compared with the control group (aCSF) (p=0.006 and p=0.042, respectively), indicating the expression of CPP. The animals receiving 70 nmol of ethanol showed the highest preference (200 ± 26 seconds vs 145 ± 20 seconds for the 150 nmol group). Interestingly, while the lowest dose administered (35 nmol) did not induce any preference, the highest dose (300 nmol) induced a significant decrease in the preference score relative to the control group (p=0.023), demonstrating the expression of CPA for this dose. The score for this dose was also significantly lower compared with that observed for animals treated with 70 nmol dose (p< 0.001). As expected, the preference score from the unpaired group conditioned with the dose that was shown to produce the highest preference (70 nmol) did not differ from the control group (aCSF) (p=1.000).

### Increase GluN2A mRNA but not GluN1 mRNA levels in NAc is associated with the expression of ethanol place preference

First, western blot technique was selected to analyse changes in NMDAR GluN1 and GluN2A subunits in the NAc and dorsal Hippocampus (Figures 1D and 1E, respectively). The analysis was performed on brain tissue extracted from the groups that showed the highest preference or aversion (70 nmol and 300 nmol) and also in the unpaired and control group. In the NAc, the percentage of change of the expression of GluN1 was not significantly different from control (F(3,12)=2.050; p=0.161) although levels for the group conditioned with 70 nmol of ethanol were higher (143 ± 12 %, expressed as percentage of change from control). Also, this same group (70 nmol treated) showed the highest level of GluN2A (166 ± 27 %, expressed as percentage of change from control); however, the Kruskal-Wallis showed no significant differences between groups (p=0.126). In the Hippocampus, GluN1 and GluN2A showed a slight tendency to be less expressed in the Hippocampus of the rats that received 300 nmol ethanol dose. In any case, the one-way ANOVA showed that the level of expression of GluN1 and GluN2A between groups was not significantly different (F(3,11)=2.078; p=0.131 and F(3,12)=1.066; p=0.403, respectively).

To further analyze the tendency in the changes of GluN2A expression, a different cohort of animals was conditioned with 70 nmol and aCSF as control group (Figure 2A). The objective was to replicate the previous behavioral results and perform a more precise analysis of the GluN1 and GluN2A expression. As shown in the previous experiment, 70 nmol of ethanol significantly increased the time spent in the drug-paired compartment (p=0.009, T-test). The *In Situ* Hybridization results (Figure 2D and 2E) revealed that, in the NAc, the group that expressed CPP after ethanol treatment, has significantly higher mRNA levels of GluN2A (5.64 ± 0.54 dots/cell) as compared with the control group (3.62 ± 0.03 dots/cell) (p=0.014, t-test). However, the T-test did not reveal any significant difference in GluN1 mRNA levels between the two groups (p=0.316). Besides, in the Hippocampus, the levels of GluN1 and GluN2A mRNAs were not different between ethanol conditioned and control animals, as shown by the T-test (p=0.896 and p=0.698). Representative pictures of the groups analysed are shown Figures 3, 4 and 5.

**Figure 4.**
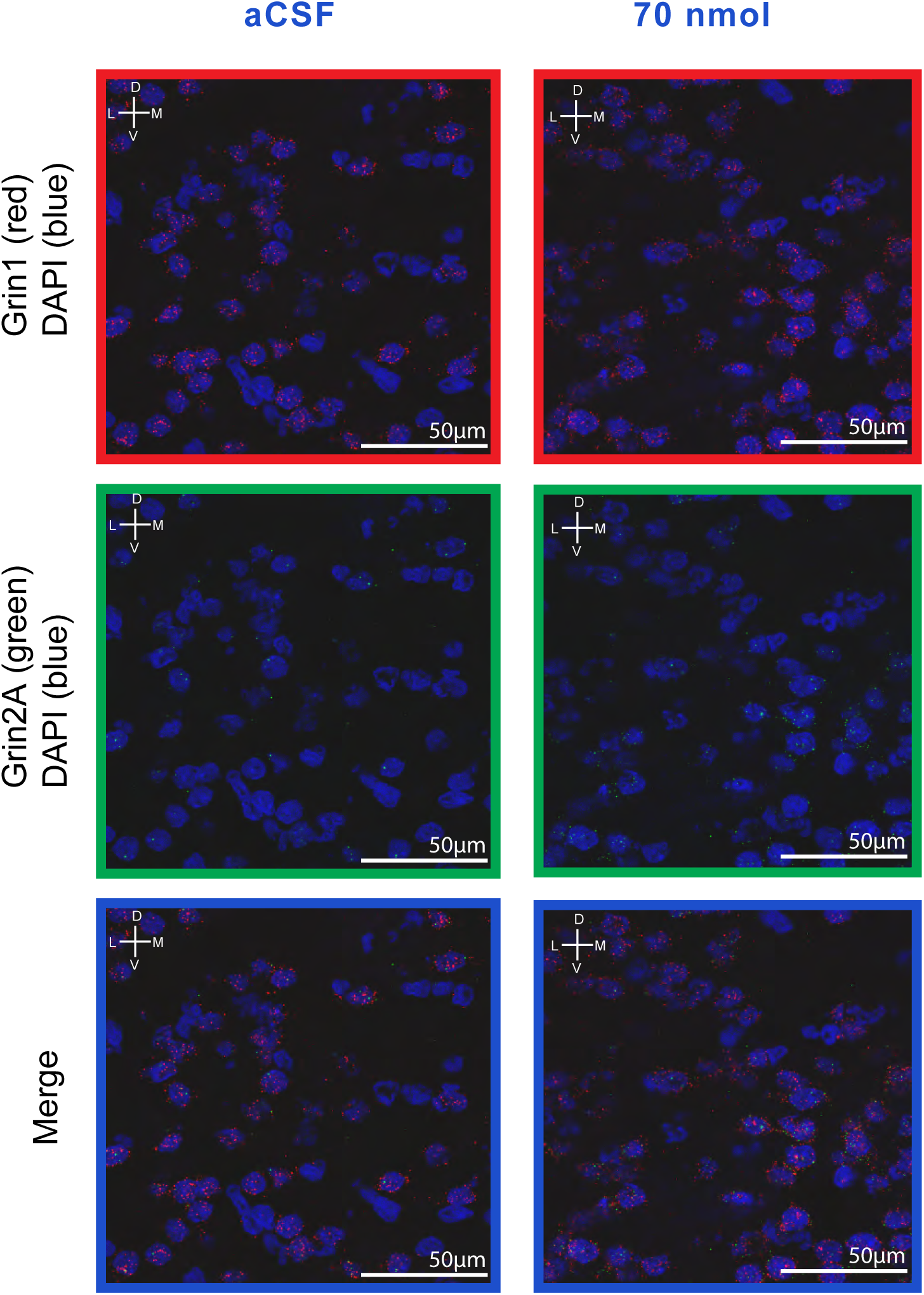
Representative amplified pictures obtained in the mRNA *In Situ* Hybridization Assay used for the quantification of mRNA from GluN1 and GluN2A subunits in NAc. Amplified pictures from one animal of each group (aCSF on the left and ethanol 70 nmol on the right) were selected.

**Figure 5.**
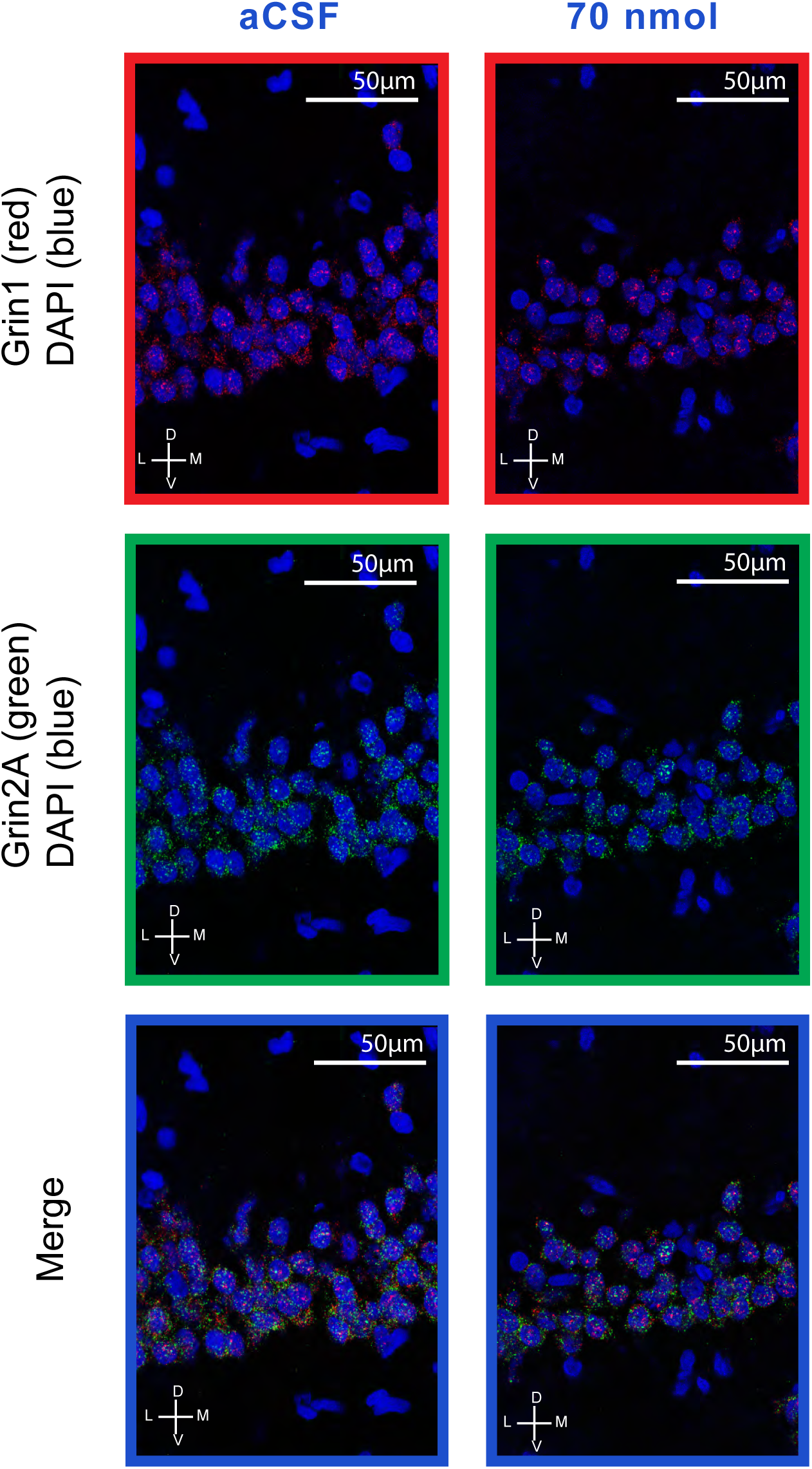
Representative amplified pictures obtained in the mRNA *In Situ* Hybridization Assay used for the quantification of mRNA from GluN1 and GluN2A subunits in hippocampus. Amplified pictures from one animal of each group (aCSF on the left and ethanol 70 nmol on the right) were selected.

### The blockade of pVTA MORs impairs the acquisition of ethanol-induced CPP

As shown in Figure 6B, the pre-treatment with β-FNA was able to block the acquisition of the CPP induced by the ethanol dose that has previously been shown to trigger the highest preference score (70 nmol). Therefore, the statistical analysis showed that only the group pre-treated with aCSF and conditioned with 70 nmol of ethanol presented a significantly higher score as compared to the three other groups (Kruskal-Wallis, p=0.002; pairwise comparisons with Bonferroni adjustment, p<0.005). Besides, using western blot, we analyzed the changes in NMDA GluN1 and GluN2A subunits expression in the NAc from the group pre-treated with β-FNA after conditioning with 70 nmol of ethanol (Figure 6D). In this case, GluN1 and GluN2A protein levels were not significantly different from those of the control group (aCSF + aCSF) as shown by the t-test and the non-parametric test (p=0.72 and p=0.670, respectively). Indeed, no tendency could be observed in these results.

**Figure 6.**
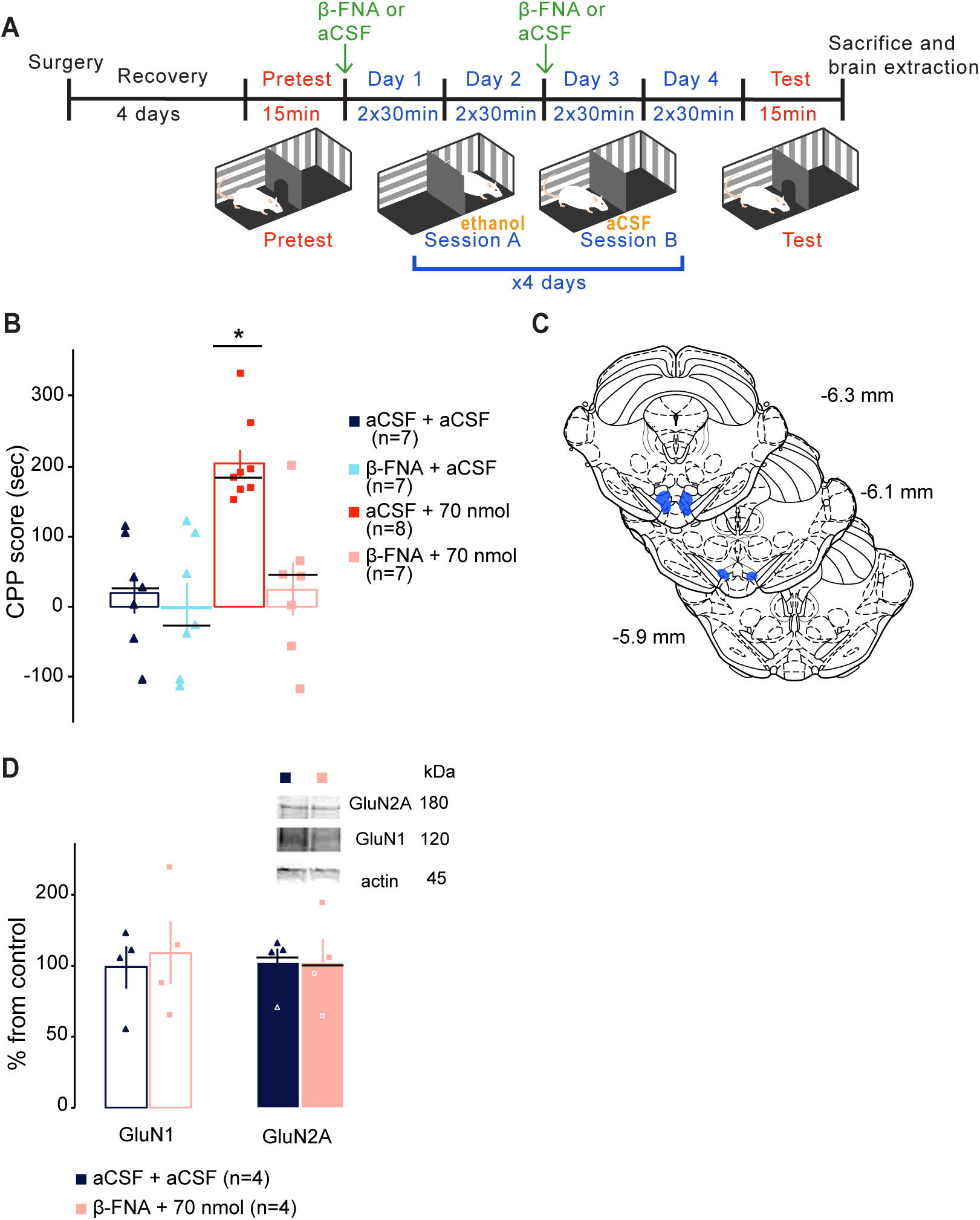
The blockade of pVTA MORs impairs the acquisition of ethanol-induced CPP. ***A***, Schematic of the experimental design. ***B***, Blockade of ethanol 70 nmol induced CPP after the pretreatment with β-FNA. Data are mean ± SEM represented as preference score (test minus pretest time spent in ethanol-paired compartment). The black line represents the median for the non-normal distributed data. * denotes significant differences relative to the control group (aCSF) (p<0.005; Kruskal-Wallis followed by Bonferroni adjustment for pairwise comparisons). ***C***, Diagram of brain coronal sections indicating in blue the area of all microinjections in pVTA. ***D***, NMDA subunit expression in NAc after pretreatment with β-FNA. Data are mean ± SEM represented as percentage from control group (aCSF + aCSF). The black line represents the median for the non-normal distributed data.

## 4. Discussion

Associations between the drug and the environment play a relevant role in the development and maintenance of Alcohol Use Disorders. In the present study, we demonstrated that the association between the ethanol local action in the pVTA and a context is able to induce CPP or CPA in an elegant dose-response manner. Furthermore, we showed here that the development of an ethanol-context preference necessarily involves the activation of the MORs within the pVTA and modifies the mRNA levels of GluN2A in the NAc, what may impact the expression of NMDAR GluN2A subunits in this structure, although no direct evidence was observed.

The ability of ethanol to induce CPP in different strains of rats has been previously described when administered systemically (Bahi and Dreyer, 2013; Ciccocioppo et al., 1999; Peana et al., 2008). In fact, studies using medium doses of ethanol such as 1 g/Kg i.p. or intragastric (gavage) and similar experimental design to the one used in this study, have reported ethanol-place preference (Bahi and Dreyer, 2013; Peana et al., 2008). Furthermore, Walker and Ettenberg showed that intracerebroventricular (icv) ethanol administration was able to induce CPP at the dose of 180 nmol whereas 120 or 240 nmol did not induce changes in the preference (Walker and Ettenberg, 2007). It is well-established that the reinforcing properties of drugs which trigger preference for the drug-associated environment are mediated by an increased activity of the VTA DA neurons. None of these studies have established a correlation between the ethanol systemic doses and brain ethanol concentrations achieved. In some of these studies, ethanol plasmatic levels were measured, however, these concentrations did not represent the actual concentration in discrete brain areas (i.e. VTA), which may be very difficult to measure whereas animals are following CPP protocol. However, to our knowledge, none of the studies using the CPP paradigm has administered ethanol locally into the pVTA. Therefore, our results show for the first time that the local intra-pVTA administration of 70 and 150 nmol of ethanol paired with a specific context results in a preference for the drug-associated environment. In line with our previous results (Martí-Prats 2013, 2015 and Hipolito 2011), we hypothesize that the products derived from the ethanol metabolism (i.e. salsolinol) could be responsible for the induction of the place preference. Figure 7 represents a schematic illustration of our hypothesis about the ethanol-induced CPP; however, some aspects of this hypothesis remain to be confirmed and clarified in future investigations. According to our hypothesis presented here, the ethanol metabolic fraction is necessary for the development of CPP. In fact, previous published work showed that the blockade of ethanol metabolism by inhibiting acetaldehyde dehydrogenase (an enzyme necessary for ethanol oxidation) or by using strategies aimed at sequestering the first ethanol metabolite, therefore preventing the formation of salsolinol, suppresses the preference induced by the systemic administration of ethanol (Peana et al., 2008). Moreover, local intra-VTA administration of salsolinol, an ethanol derivative, is able to increase DA extracellular levels in the NAc and induce CPP (Hipólito et al., 2011). Thus, ethanol derivatives in general and salsolinol in particular, could be responsible for the ethanol-induced activation of the VTA dopaminergic neurons which could lead to an increase of DA release in the NAc, a mechanism necessary to induce context-learned associations (Hipólito et al., 2011). This activation could be mediated by the action of salsolinol over MORs and not by the ethanol molecule itself. In line with recent studies showing that acquisition of ethanol-induced CPP (i.p. or icv) can be prevented by inhibiting MORs (Gajbhiye et al., 2017; Gibula-Bruzda et al., 2015; Quintanilla et al., 2014), we showed here that the blockade of local pVTA MORs during the conditioning phase impaired the acquisition of ethanol place preference. This highlights the key role of the MORs in the VTA in the development of ethanol induced context-learned associations. Moreover, recent studies have demonstrated that salsolinol is able to act as an agonist of the MORs (Berríos-Cárcamo et al., 2016; Hipólito et al., 2011; Quintanilla et al., 2014; Xie et al., 2012). To our knowledge, there is no evidence in the literature of direct ethanol interaction with the MORs (Melis et al., 2015) although it is clearly stated that MORs antagonists are at some level useful to reduce alcohol compulsive intake (Soyka et al., 2016).

**Figure 7.**
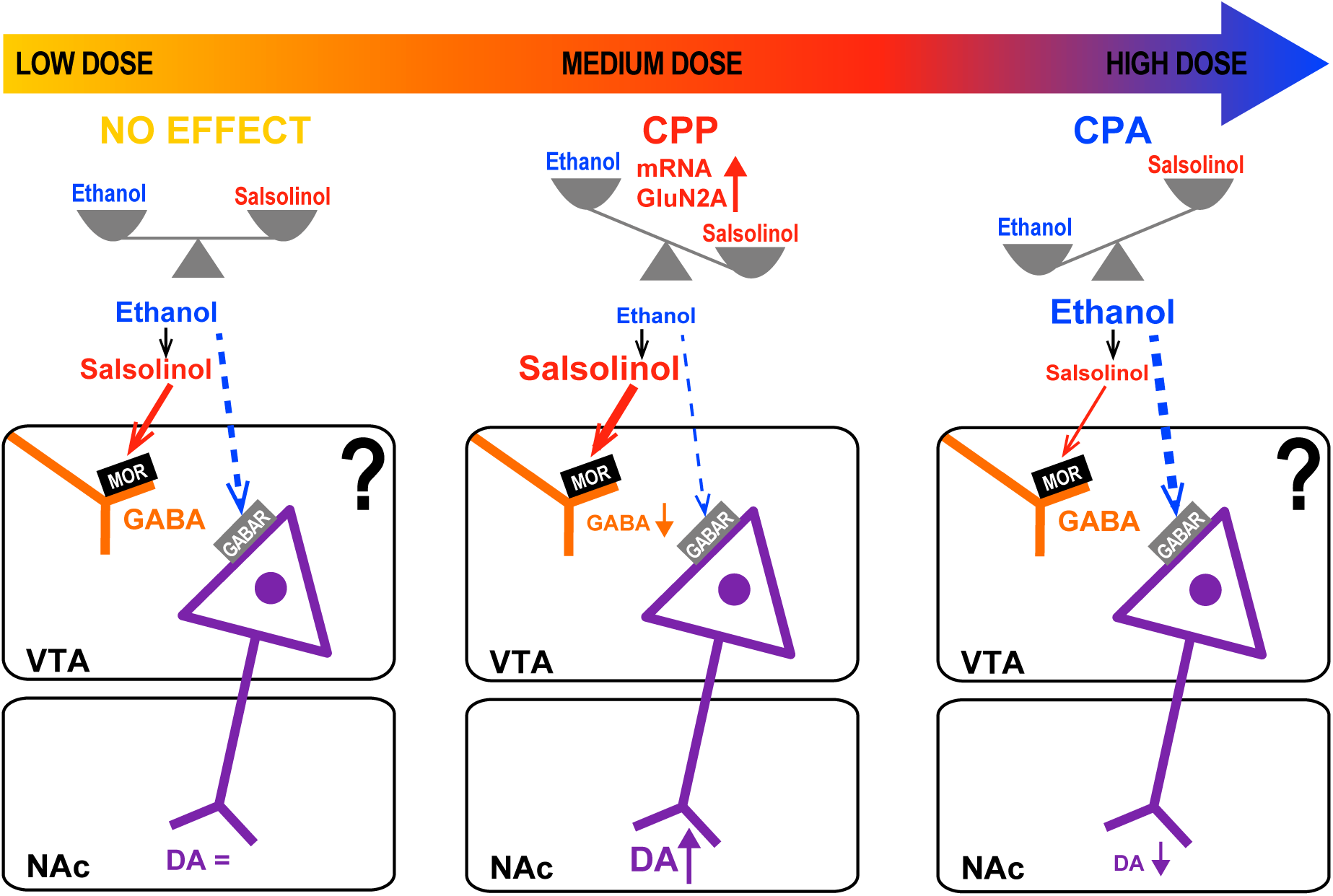
Simplified representation of the dose-dependent ethanol effect on place preference or place aversion and its hypothesized mechanism. The administration of low doses of ethanol into the VTA does not induce preference or aversion for the drug paired compartment. Medium doses elicit CPP, presumably through an increase of the ethanol metabolized fraction. In our hypothesis and based in previous literature, the ethanol metabolite, salsolinol, would activate MORs, decreasing GABA release and finally increasing DA release in the NAc by a disinhibition of VTA DA neurons. The expression of ethanol induced CPP also results in an increase of mRNA form GluN2A in the NAc. Moreover, the blockade of the MORs in the VTA inhibits the acquisition of this ethanol induced CPP. However, the administration of high doses results in the development of CPA. In this case, the effect of the non-metabolized fraction would predominate and the inhibition of the VTA DA neurons would result in a decrease of DA release in the NAc. In orange GABA terminals and in purple VTA DA neurons. MOR: mu opioid receptor; GABAR: gamma-amino butyric acid (GABA) receptor; NAc: nucleus accumbens; VTA: ventral tegmental area; DA: dopamine; GLUN2A: NMDA receptor subunit 2A.

Interestingly, ethanol administration not only can induce CPP but also can result in an aversion to the drug-paired compartment when a higher dose is administered. This is observed in our study when animals are treated with 300 nmol ethanol and show a clear negative preference score. There is also previous evidence of ethanol-induced CPA in the literature, particularly when high systemic ethanol doses (2 g/kg) were administered (Becker et al., 2006). According to our hypothesis, this ethanol-induced place aversion could be a consequence of the direct action of the ethanol molecule itself (*i.e.* the non-metabolized fraction of the administered dose) in the VTA (Figure 7). Although no direct evidence of the aversion mechanism is investigated in this current study, larger doses may result in accumulation of ethanol which has been demonstrated to exert depressant effects on VTA DA-neurons (Martí-Prats et al., 2013; Theile et al., 2011, 2009, 2008; Xiao and Ye, 2008) and might induce CPA (Tan et al., 2012). These data, altogether with our and other previous data, clearly indicate that understanding the mechanism of action of ethanol in the VTA underlying the expression of reinforcement or aversion is still unknown and more efforts are needed to shed light on this important issue, especially to understand ethanol-mediated aversion mechanisms.

Other neurotransmitter systems are also important in mediating the drug-elicited context dependent associations. Indeed, the development and expression of these drug-context learning associations has been classically associated with glutamatergic transmission and glutamatergic-dependent neural plasticity induced by drugs (Hearing et al., 2018). In the present study, increased GluN2A mRNA dots/cell counts, together with a tendentious increase in GluN2A protein levels in the NAc after ethanol place preference were observed. It is remarkable that these changes are attributable to the establishment of an association between the ethanol effects and a specific drug-conditioning compartment because ethanol-unpaired rats exhibit no changes in NMDAR expression. Thus, it could be possible that accumbal GluN2A play a key role in the development of the association between ethanol and the environment. In fact, GluN2A knockout mice are not able to develop morphine or ethanol CPP (Boyce-Rustay and Holmes, 2006; Miyamoto et al., 2004), whereas blockade of GluN2B does not alter ethanol place preference (Boyce-Rustay and Cunningham, 2004). In addition, local antagonism of NMDA receptors in the NAc before the test session blocks the expression of ethanol-induced CPP (Gremel and Cunningham, 2009). However, we are still not able to discern if the changes in GluN2A mRNA reflect an increased expression of functional GluN2A subunits in the NAc and moreover if these changes play a role in the ethanol induced CPP expression. Several aspects, from sample size and limitations of the western blot technique to post-transcriptional mechanisms, may explain why we could not observe changes in the functional protein expression. Clearly, further studies would be needed to explore this idea and finally state if increases observed in GluN2A mRNA levels translate into more expression of GluN2A subunit. Interestingly and in line with our results in the Hippocampus, ethanol place preference does not seem to induce changes in the expression of GluN1 nor GluN2A subunits in this region. This is an intriguing result because previous studies have shown that morphine CPP in mice is related to an increase of GluN1 subunit in the Hippocampus (Portugal et al., 2014). In addition, the local antagonism of NMDA receptors in the Hippocampus has also be shown to prevent the acquisition but not the expression of morphine-induced place preference (Zarrindast et al., 2007). Importantly, in these experiments, morphine was administered systemically in contrast to our current experiments whereby ethanol was locally administered into the pVTA. Unfortunately, to the best of our knowledge, there are no studies that analyzed the expression of these NMDA subunits in the Hippocampus after the development of ethanol CPP. It could then be possible that the adaptations in the Hippocampus potentially occurring as a consequence of the association of ethanol with a specific context, could differ from those elicited by other opioidergic drugs or may be related to the drug-induced activation of MORs located in other brain areas. Further investigation may help understanding the role of Hippocampal NMDAR in the expression of ethanol-induced CPP.

## 5. Conclusion

Altogether, these data offer new avenues to further our knowledge about the mechanism of action of ethanol in the VTA and its associated neurochemical and behavioral effects. The opposite effect exerted by low versus high doses of ethanol triggering variety of results in the literature has ultimately revealed the complexity of ethanol action in the MCLS. Our results shed some light on the mechanism underlying the ethanol-induced context learned associations, the MORs and the ethanol selected dose being crucial in the development of the CPP. Additionally, the acquisition of CPP might induce adaptations in the NMDA subunit composition in the NAc originating from the ethanol action on the VTA. As suggested by our *in situ* hybridization experiments, GluN2A mRNA increase may play a role in the ethanol-induced plasticity that facilitates the expression of the CPP. These novel results set the need for further analysis on the mechanism by which ethanol develops and expresses its motivational properties associated with learned environmental cues.

## 6. Acknowledgements

This work was supported by University of Valencia “Special grants” UV-INV-AE13-136776 (L.G.); Spanish Ministerio de Economia y Competitividad’’ MINECO PSI2016-77895-R (L.H.); US National Institutes of Health (NIH) grant DA041781 (J.A.M.), DA042581 (J.A.M.), DA042499 (J.A.M.), DA041883 (J.A.M.), NARSAD Independent Investigator Award from the Brain and Behavior Research Foundation (J.A.M.). We would like to thank the students Ms. Valentina Nurra and Mr. Andrés Martínez-Pardo participating in some of the experiments and the lab technician Mr. Jose Luís González-Romero. We also thank Ms. Pilar Laso for grant management and personnel of the Animal facilities of the SCSIE, University of Valencia, for assuring animal welfare.

## 7. Authors contribution

Conceptualization, L.M.P, A.P., L.G., L.H.; Methodology, Y.C.J., L.M.P., J.A.M., A.P., L.G., L.H; Formal Analysis, Y.C.J., L.M.P, A.P., L.G, L.H; Investigation, Y.C.J., L.M.P Writing – Original Draft, Y.C.J, L.H; Resources, J.A.M., A.P., L.G., L.H; Supervision, L.M.P, J.A.M., A.P., L.G., L.H. Writing – Review & Editing, Y.C.J., L.M.P., J.A.M., A.P., L.G., L.H; Funding Acquisition, J.A.M, L.H., L.G., A.P.

